# AlphaMissense pathogenicity scores predict response to immunotherapy and enhances the predictive capability of tumor mutation burden

**DOI:** 10.1101/2025.09.28.679078

**Authors:** David Adeleke, Adewale Oluwaseun Fadaka, Nicole Remaliah Samantha Sibuyi, Ashwil Klein, Mervin Meyer, Gomes Rahul, Rick Jansen

## Abstract

Tumor Mutational Burden (TMB) is a widely used biomarker for selecting cancer patients for immune checkpoint inhibitor (ICI) therapy. However, TMB alone has limited predictive power, as it fails to account for the functional impact of mutations. We introduce AlphaTMB, a composite biomarker that integrates the quantity of mutations (TMB) with the qualitative assessment of their pathogenicity using AlphaMissense, a deep learning model that predicts the deleteriousness of missense variants. Using a pan-cancer cohort of 1,662 patients from the MSK-IMPACT study who received ICI therapy, we computed three scores per patient: TMB, Alpha (sum of AlphaMissense scores), and AlphaTMB (product of TMB and Alpha). Patients were stratified using both cancer-specific and pan-cancer quantiles. Survival outcomes were evaluated using Kaplan-Meier and multivariate Cox proportional hazards models, controlling for cancer type, age, and ICI regimen. AlphaTMB showed strong correlation with TMB (Spearman ρ = 0.866, *p* < 0.001), but offered improved prognostic accuracy. Patients in the bottom 80% AlphaTMB group had significantly poorer survival than those in the top 10% (HR < 2.51, *p* < 0.001), outperforming TMB and Alpha alone. AlphaTMB reclassified borderline cases, identifying subsets with low TMB but high deleterious mutation load, and vice versa. Gene mutation heatmaps and co-occurrence analysis confirmed that to 10% AlphaTMB-high tumors were enriched in mismatch repair and POLE mutations, reflecting a neoantigen-rich, immunotherapy-responsive phenotype. AlphaTMB improves survival prediction beyond TMB alone, better captures immunogenic tumor profiles, and reflects more accurate patient stratification. This AI derived somatic mutations pathogenicity scoring represents a step toward personalized immuno-oncology and merits further validation in prospective studies.

## 1. Introduction

Cancer remains a major global health challenge, accounting for nearly 20 million new cases and 9.7 million deaths in 2022 alone, with incidence and mortality rates projected to rise significantly over the coming decades [1]. Although advances in early detection and targeted therapies have improved outcomes in high-income countries, global disparities in access to care persist, especially in low- and middle-income countries where late-stage diagnoses and limited treatment access contribute to poorer outcomes [2–4].

In recent years, immune checkpoint inhibitors (ICIs) have revolutionized cancer therapy by enabling durable responses in a subset of patients across a broad spectrum of malignancies, including non-small cell lung cancer (NSCLC), melanoma, bladder cancer, and renal cell carcinoma [5–8]. These agents, which target immune inhibitory pathways such as PD-1/PD-L1 and CTLA-4, restore the ability of cytotoxic T cells to recognize and eliminate tumor cells. However, despite their success, only a minority of patients derive substantial clinical benefit, and response rates remain highly variable [9]. This variability underscores an urgent need for robust biomarkers to predict which patients are most likely to benefit from ICI therapy.

Tumor Mutational Burden (TMB), defined as the total number of nonsynonymous somatic mutations per megabase of tumor DNA, has emerged as a promising biomarker for immunotherapy response. The underlying premise is that tumors with a high mutational burden are more likely to generate neoantigens capable of eliciting an effective immune response [10, 11]. Retrospective studies have shown that TMB correlates with objective response rate and progression-free survival in patients treated with ICIs, particularly in tumors such as melanoma, NSCLC, and urothelial carcinoma [12, 13]. These observations culminated in the U.S. Food and Drug Administration’s tissue-agnostic approval of pembrolizumab for solid tumors with TMB ≥10 mutations/Mb, based on findings from the KEYNOTE-158 trial [14].

However, the clinical utility of TMB has been increasingly called into question. First, TMB does not capture the biological significance or immunogenicity of individual mutations. It is a purely quantitative measure. Not all mutations contribute equally to neoantigen formation or immune activation; many are biologically neutral passenger mutations with minimal relevance to cancer progression or immune escape [15–18]. Second, the threshold defining “TMB-high” is not standardized and may vary significantly across tumor types, sequencing platforms, and gene panel sizes [19, 20]. Third, tumors with similar TMB levels often exhibit divergent clinical responses, suggesting that the quality not just the quantity of mutations may be essential for predicting therapeutic benefit [21, 22].

To address these limitations, recent efforts have focused on refining mutational biomarkers by incorporating measures of functional impact. In this context, machine learning-based algorithms offer new opportunities to predict the pathogenicity of individual mutations. AlphaMissense, a deep learning model developed by Google DeepMind, predicts the deleteriousness of missense mutations by integrating protein structural data from AlphaFold, evolutionary conservation, and sequence context [23]. Unlike earlier tools such as SIFT or PolyPhen-2, AlphaMissense achieves superior accuracy in differentiating benign from pathogenic variants, making it an ideal candidate for refining existing genomic biomarkers. Building on this concept, we propose AlphaTMB, a composite metric that combines the mutation count (TMB) with the predicted pathogenicity of each variant, as determined by AlphaMissense. This integrated score aims to distinguish tumors harboring a high load of biologically meaningful, potentially immunogenic mutations from those with a high burden of functionally neutral variants. We hypothesize that AlphaTMB will offer improved prognostic performance over TMB alone, enabling more precise stratification of patients for immunotherapy.

To test this hypothesis, we leveraged data from the MSK-IMPACT cohort a large, real-world clinical sequencing dataset of over 1,600 patients treated with ICIs across multiple cancer types. By comparing TMB, AlphaMissense scores, and the integrated AlphaTMB score, we assessed their respective associations with overall survival and evaluated their performance in patient classification across global and cancer-specific contexts. Our findings suggest that incorporating functional mutation scoring into TMB provides a more refined and biologically meaningful biomarker that better predicts patient outcomes on immunotherapy.

This study contributes to the growing body of evidence advocating for multi-parametric, functionally informed biomarkers in oncology. As precision cancer medicine moves beyond one-size-fits-all models, tools like AlphaTMB hold promise for enhancing therapeutic decision-making and optimizing patient benefit from immunotherapy.

## 2. Methods

### 2.1 Study Cohort and Data Acquisition

This study analyzed genomic and clinical data from the MSK-IMPACT (Integrated Mutation Profiling of Actionable Cancer Targets) cohort, a large-scale sequencing initiative conducted at Memorial Sloan Kettering Cancer Center and publicly available via the cBioPortal for Cancer Genomics. We selected a subset of 1,662 patients who had undergone next-generation sequencing using one of the MSK-IMPACT gene panels and had received at least one dose of immune checkpoint inhibitor (ICI) therapy. Treatment regimens included anti–PD-1, anti– PD-L1, and anti–CTLA-4 monotherapies or combinations, specifically involving agents such as atezolizumab, avelumab, durvalumab, ipilimumab, nivolumab, pembrolizumab, and tremelimumab. Clinical data, including tumor type, age, sex, ICI regimen, and overall survival status, were retrieved using the cBioPortal R API. For patients who received more than one ICI-based treatment, only the first documented regimen was included in the analysis. Survival was measured from the date of initial ICI administration, and patients were censored at the date of last follow-up if death was not recorded. **Available covariates and data constraints:** The MSK-IMPACT ICI dataset included age, sex, tumor type, immune checkpoint inhibitor (ICI) class, and MSK-IMPACT panel version (341/410/468). ECOG performance status, prior therapy history, tumor stage, PD-L1 expression, MSI/dMMR status, HLA genotype, and tumor microenvironment features were not available in this dataset [24].

### 2.2 Computation of Mutation Burden and Pathogenicity Scores

Three mutational metrics were computed for each tumor sample. Tumor Mutational Burden (TMB) was defined as the total number of nonsynonymous somatic mutations per megabase of sequenced exonic DNA, normalized to the coverage size of the corresponding MSK-IMPACT panel (0.9 Mb for IMPACT341, 1.1 Mb for IMPACT410, and 1.3 Mb for IMPACT468). Panel version was included as a covariate in multivariable models to mitigate residual platform effects. In parallel, an AlphaMissense pathogenicity score was calculated by summing the predicted deleteriousness scores of all missense mutations in each sample, as determined by DeepMind’s AlphaMissense model. This model leverages evolutionary conservation and AlphaFold-derived structural information to predict the functional impact of amino acid substitutions. To create a composite measure reflecting both mutation load and functional significance, we defined AlphaTMB as the product of TMB and the total AlphaMissense score. All metrics were standardized across the cohort using z-score transformation to ensure comparability. Immune-relevant genes refer to literature-curated genes involved in antigen presentation or immune evasion; no predictive modeling was performed.

### 2.3 Patient Stratification by Mutation Metrics

Patients were stratified into risk groups using two percentile-based methods. In the global percentile approach, thresholds for the Top 10%, Top 10–20%, and Bottom 80% groups were calculated across the entire cohort, independent of tumor type. This facilitated pan-cancer comparisons of mutational biomarkers. In the cancer-type–specific approach, percentiles were calculated within each tumor type, allowing classification relative to the mutational landscape of that specific cancer. Both approaches were applied independently to the TMB, AlphaMissense, and AlphaTMB metrics to explore their performance in different clinical contexts. Percentile-based stratification (Top 10%, Top 10–20%, Bottom 80%) was used to avoid imposing linearity assumptions and to align with prior TMB literature, enabling pan-cancer comparability.

### 2.4 Survival Analysis and Statistical Modeling

Overall survival (OS) was evaluated using the Kaplan-Meier method, and statistical differences between groups were assessed using the log-rank test. To control for potential confounding factors, multivariable Cox proportional hazards regression models were constructed, incorporating age, tumor type, and ICI treatment class as covariates. For each cancer type, cases were stratified into the top 20% percentile for each metric, and the log-rank p-value was calculated to assess differences in overall survival (OS). The direction of the effect was evaluated, and a hazard ratio (HR) was determined using a Cox proportional hazards model. Additional analyses were performed with 10% to 50% metric cutoffs. Hazard ratios (HRs) and 95% confidence intervals (CIs) were calculated to estimate the relative risk of death associated with each mutational risk group. **Drug-class analyses:** Survival analyses were repeated by ICI class (anti– PD-(L)1, anti–CTLA-4, and combination regimens) to explore whether biomarker performance varied by treatment modality. All statistical analyses were performed using R version 4.2.2, with visualization and modeling conducted using the “survival” and “survminer” packages. A bootstrap resampling framework (1,000 iterations) is proposed for future work to assess model stability and optimism-corrected c-indices.

## 3. Result

To evaluate the clinical and biological relevance of mutation burden and functional impact in immunotherapy-treated cancers, we systematically analyzed Tumor Mutational Burden (TMB), AlphaMissense scores, and the composite AlphaTMB metric across a pan-cancer cohort of 1,662 patients. We began by characterizing the distribution and correlation of these genomic scores, followed by assessments of their prognostic value using survival analyses. Across ICI classes, AlphaTMB maintained prognostic discrimination similar to TMB in anti–PD-(L)1 subsets and combination therapy, with wider confidence intervals in smaller strata. We then examined the ability of AlphaTMB to refine patient classification beyond existing TMB- and Alpha-based groupings. Finally, we investigated mutation-level patterns within each scoring framework to determine whether AlphaTMB more accurately captures immunogenic tumor phenotypes associated with favorable outcomes. Together, these analyses provide a comprehensive view of how integrating mutation quantity with predicted functional impact enhances the predictive resolution of existing biomarkers in the context of immune checkpoint blockade.

### Patient Demographics and Clinical Characteristics

Table 1 summarizes the distribution of cancer types, sex, age groups, and survival status among the 1,529 patients analyzed. Non-small cell lung cancer (NSCLC) and melanoma were the most prevalent tumor types. The majority of patients were male and aged between 50 and 70 years, reflecting the typical demographic profile of advanced cancer cases eligible for immune checkpoint inhibitor therapy.

**Table 1:**
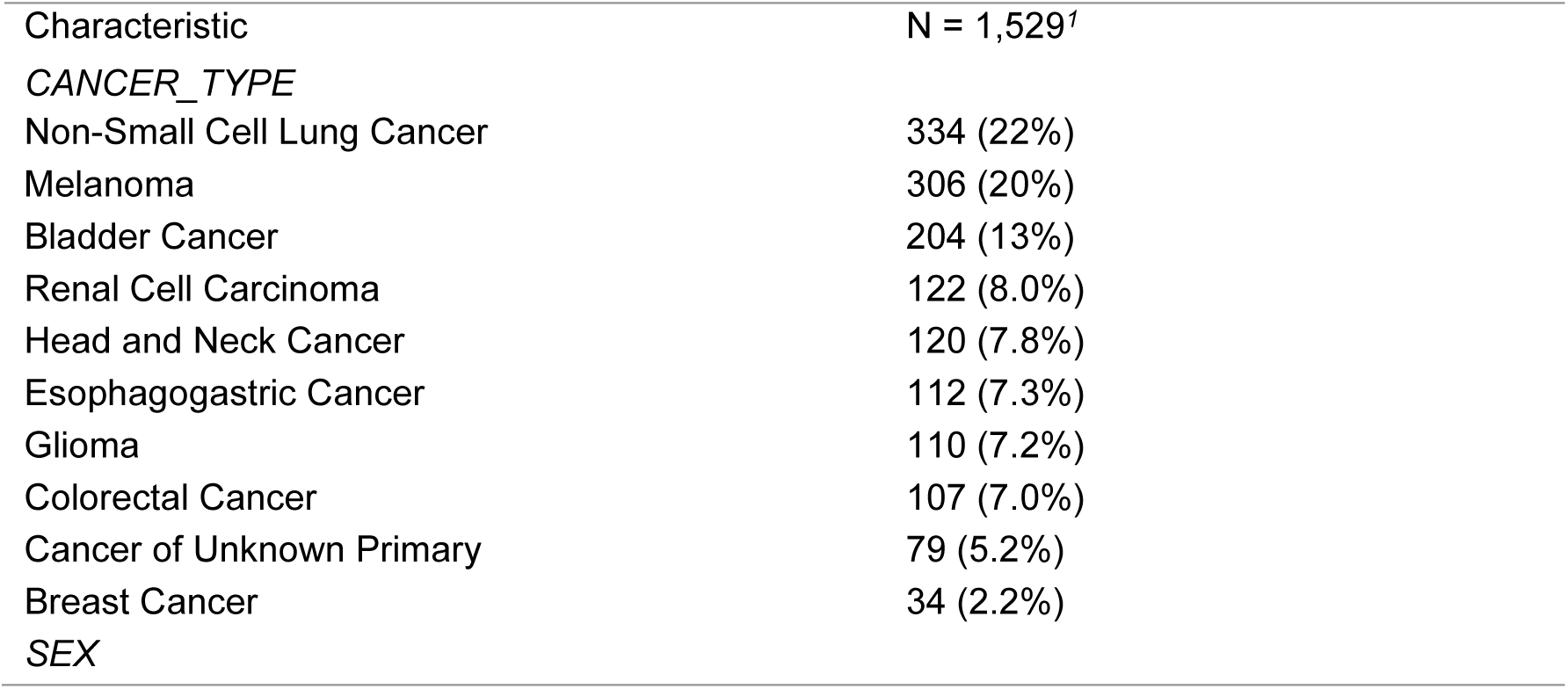
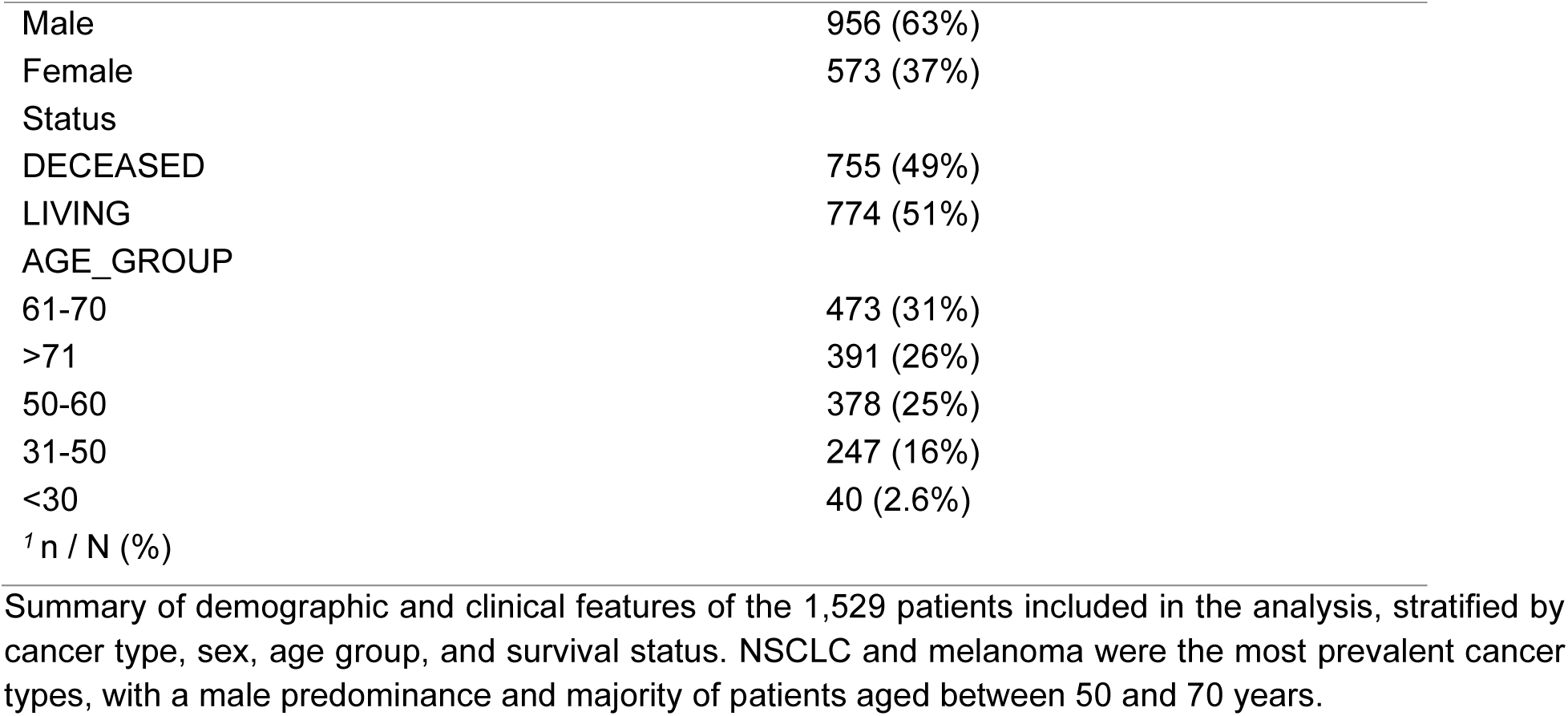
Clinicopathological Characteristics of the Immunotherapy-Treated Cohort.

| Characteristic | N = 1,529<sup>†</sup> |
| --- | --- |
| *CANCER_TYPE* |  |
| Non-Small Cell Lung Cancer | 334 (22%) |
| Melanoma | 306 (20%) |
| Bladder Cancer | 204 (13%) |
| Renal Cell Carcinoma | 122 (8.0%) |
| Head and Neck Cancer | 120 (7.8%) |
| Esophagogastric Cancer | 112 (7.3%) |
| Glioma | 110 (7.2%) |
| Colorectal Cancer | 107 (7.0%) |
| Cancer of Unknown Primary | 79 (5.2%) |
| Breast Cancer | 34 (2.2%) |
| *SEX* |  |
| Male | 956 (63%) |
| Female | 573 (37%) |
| Status |  |
| DECEASED | 755 (49%) |
| LIVING | 774 (51%) |
| AGE_GROUP |  |
| 61-70 | 473 (31%) |
| >71 | 391 (26%) |
| 50-60 | 378 (25%) |
| 31-50 | 247 (16%) |
| <30 | 40 (2.6%) |
| <sup>1</sup> n / N (%) |  |

Summary of demographic and clinical features of the 1,529 patients included in the analysis, stratified by cancer type, sex, age group, and survival status. NSCLC and melanoma were the most prevalent cancer types, with a male predominance and majority of patients aged between 50 and 70 years.

### Correlation Between Tumor Mutation Burden, AlphaMissense Scores, and Patient Survival Trends

We first examined the descriptive characteristics and relationships between TMB, AlphaMissense scores, and patient survival. As shown in Figure 1A, approximately 69.4% of tumor mutations across the cohort were successfully annotated with AlphaMissense pathogenicity scores, allowing for broad integrative analysis. This scatter plot shows the relationship between Tumor Mutation Burden (TMB), a well-established biomarker, and the AlphaMissense Score, a newly proposed metric (Figure 1B). A strong positive correlation was observed between TMB and AlphaMissense scores (Figure 1B), with a Spearman coefficient of 0.866 (*p* < 0.001), suggesting that while the two metrics are related, AlphaMissense may provide complementary functional information beyond mutation count. To explore the prognostic relevance of these scores, we analyzed their distribution according to survival status. The figure consists of two density distribution plots illustrating the distribution of Tumor Mutation Burden (Figure 1C) and AlphaMissense Score (Figure 1D), stratified by survival status (Death vs. Living). As shown in Figure 1C and 1D, both TMB and AlphaMissense scores displayed right-skewed distributions, with the majority of tumors exhibiting low to moderate values. Notably, patients who remained alive at the time of last follow-up were more likely to have higher TMB and AlphaMissense scores, whereas deceased patients were enriched among those with low mutational scores. These patterns support the association between elevated mutational burden particularly of functionally impactful variants and improved outcomes in the context of immune checkpoint blockade.

**Figure 1.**
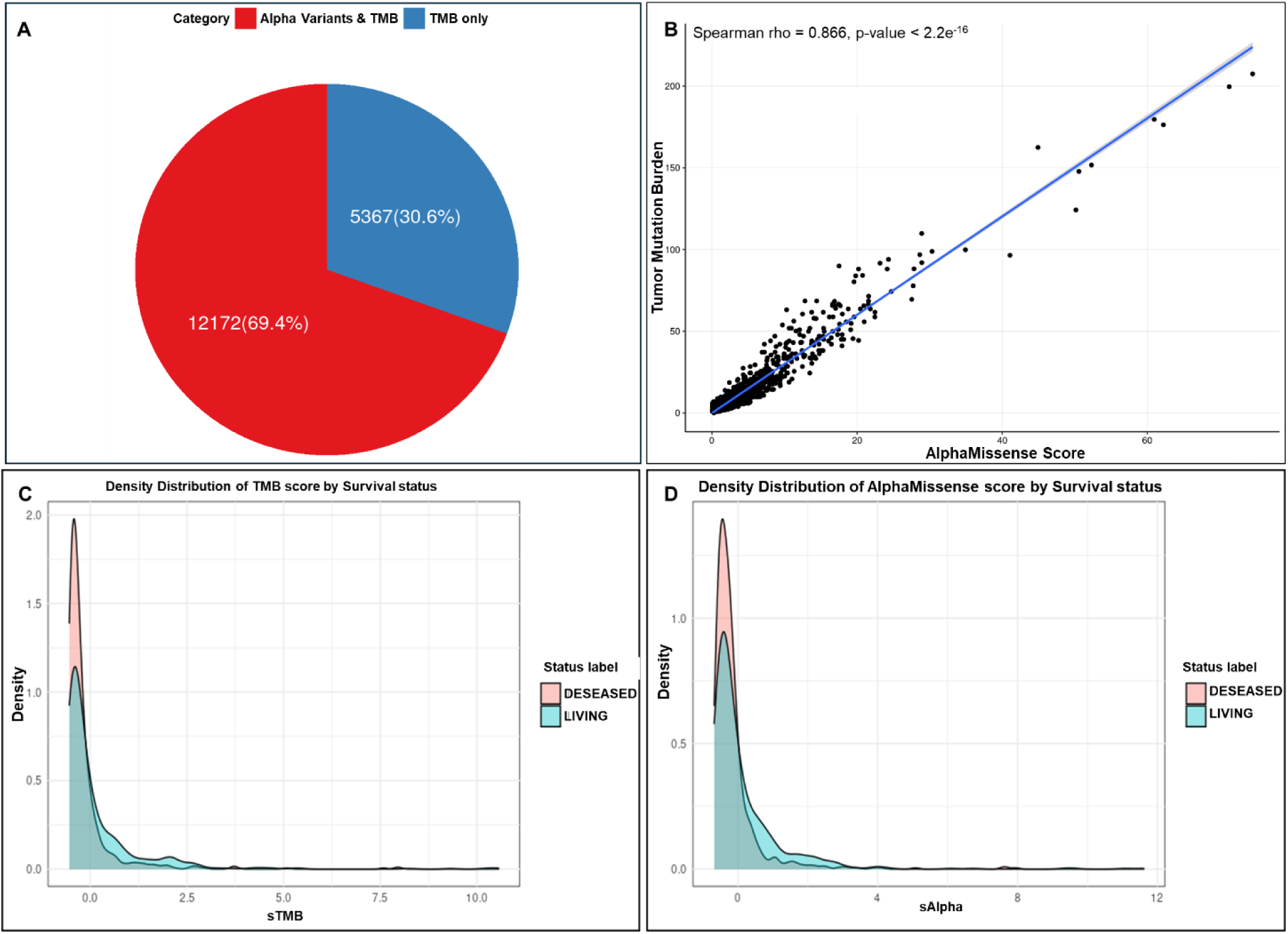
Descriptive Overview of Tumor Mutational Burden, AlphaMissense Scores, and Their Association with Patient Survival (A) Proportion of TMB variants across the cohort with an available AlphaMissense score. Approximately 69.4% of all variants were annotated with a predicted functional impact, while 30.6% lacked AlphaMissense scores. (B) Correlation plot between TMB and cumulative AlphaMissense scores per patient, showing a strong positive Spearman correlation (ρ = 0.866, *p* < 0.001), indicating that AlphaMissense captures mutation burden trends while adding functional insight. The x-axis represents the AlphaMissense **Score**, while the y-axis denotes TMB. Each data point corresponds to an individual sample, and the fitted regression line (blue) visually represents the correlation between these variables. (C) Density distribution of TMB stratified by survival status. Patients who were alive at last follow-up had a higher proportion of tumors with elevated TMB scores, while deceased patients clustered more prominently in the low-TMB range. (D) Density distribution of AlphaMissense scores stratified by survival status, revealing a similar right-skewed pattern and survival trend as TMB. Together, these panels highlight the prognostic value of both mutation burden and predicted pathogenicity.

### AlphaTMB Refines Patient Stratification Beyond TMB and AlphaMissense Alone

To evaluate the additive value of combining mutation count and pathogenicity, we examined how AlphaTMB reclassified patients previously stratified by either TMB or AlphaMissense score alone Figure 2. The two bar charts represent different regroupings based on AlphaTMB scores. The left chart shows TMB classifications reassigned using AlphaTMB scores. Similarly, the right chart does the same for Alpha-based classifications, showing how they are affected by the hybrid AlphaTMB approach. Most of the Top 10% class remains consistent (95%) in both methods, with small shifts of around 4-5% into different categories. The majority of patients within the top 10% and bottom 80% remained consistently classified under AlphaTMB, suggesting strong alignment in the most extreme risk categories. However, among patients in the Top 10–20% group, AlphaTMB led to measurable reassignments up to 5% of patients were reclassified into either higher or lower categories. These shifts were not random but reflect underlying discrepancies between mutation quantity and functional impact. For instance, tumors with modest TMB but enriched for highly deleterious mutations (high Alpha) were upgraded, whereas tumors with high mutation counts but low predicted pathogenicity were downgraded. This reclassification implies that AlphaTMB can better capture the biological relevance of the tumor’s mutational landscape, enabling better resolution of patient risk and potential immunotherapy responsiveness.

**Figure 2.**
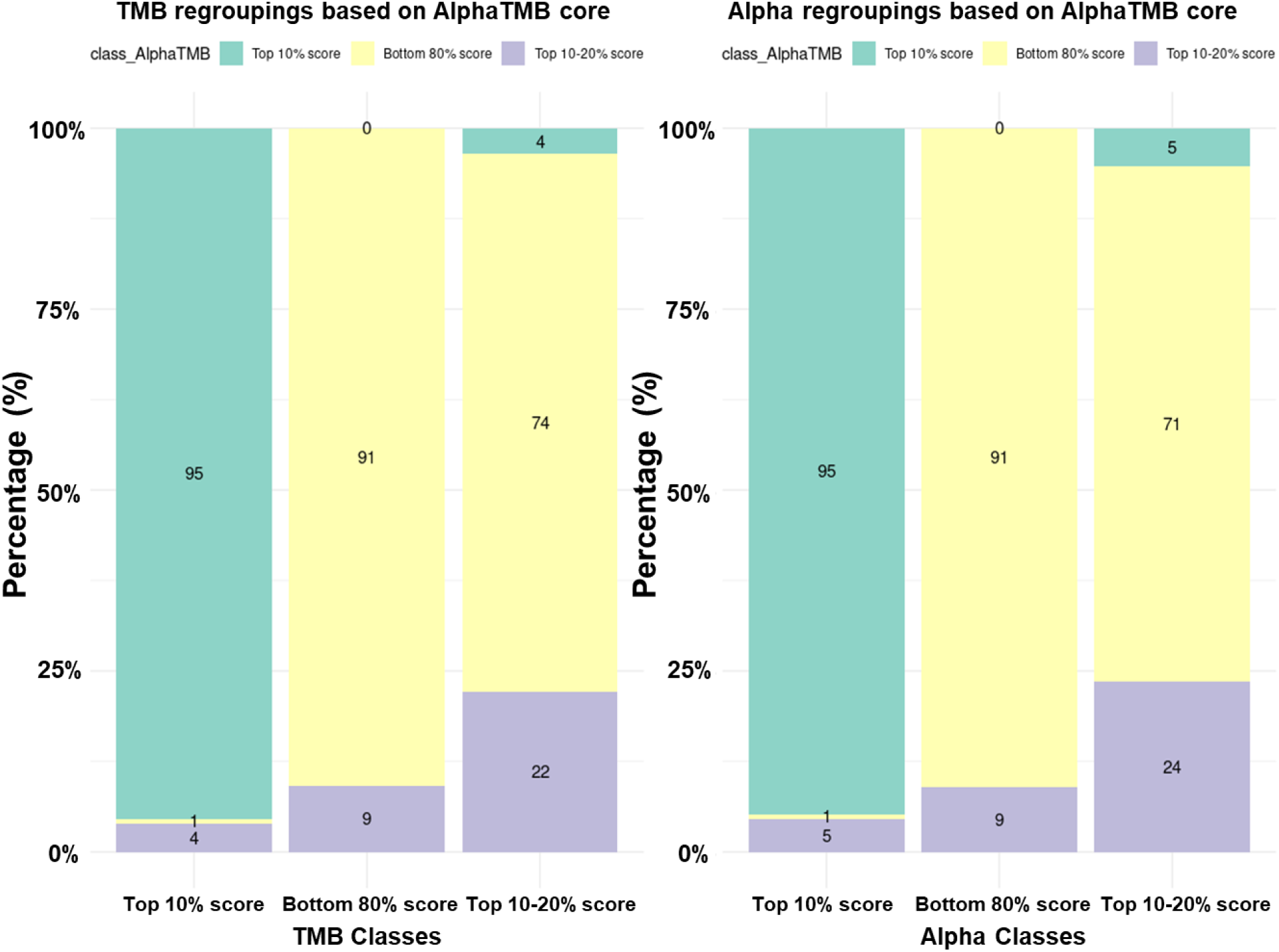
AlphaTMB Reclassifies Patient Risk Beyond TMB and AlphaMissense Alone. Bar plots showing patient redistribution when AlphaTMB is applied to reclassify TMB-based (left) and Alpha-based (right) percentile groups. The figure highlights that most patients originally classified in the Top 10% and Bottom 80% by TMB or Alpha retain their classification under AlphaTMB. However, notable reassignments occur in the Top 10–20% category, indicating that AlphaTMB captures additional functional variation not accounted for by mutation count or pathogenicity alone. These adjustments demonstrate the value of a composite metric in refining prognostic stratification.

### Pathogenicity-Weighted Mutation Burden Improves Risk Stratification and Survival Prediction in Immunotherapy-Treated Cancers

To investigate the biological differences captured by each scoring metric, we analyzed the distribution of gene-level mutations across the TMB, AlphaMissense, and AlphaTMB classifications. As shown in Figure 3A, tumors in the Top 10% TMB group exhibited broad mutational enrichment across numerous genes, including canonical drivers such as *TP53*, *KMT2D*, *TERT*, and *APC*. This pattern reflects the accumulation of mutations in hypermutated tumors but does not distinguish between functional and passenger alterations. In contrast, stratification by AlphaMissense score (Figure 3B) revealed a narrower mutation profile, with high Alpha groups showing preferential enrichment of mutations in genes associated with mismatch repair deficiency and chromatin remodeling, such as *KMT2D*, *ARID1A*, and *PTPRT*. This supports the utility of AlphaMissense in highlighting functionally impactful variants that may drive immune visibility. The AlphaTMB-based heatmap (Figure 3C) displayed the most distinct clustering of biologically relevant mutations. Samples in the top AlphaTMB group showed concentrated mutations in both known drivers and immunotherapy-relevant genes, suggesting that the combined score effectively prioritizes tumors with both high mutation load and functional immunogenicity. Compared to TMB or AlphaMissense alone, AlphaTMB more clearly delineated mutation. Of note, The heatmaps in Figure 3 illustrate mutation presence/absence and cannot capture functional weighting; therefore, they do not demonstrate statistical differences in co-occurrence patterns between metrics. AlphaTMB’s added value lies in integrating mutation quantity with predicted functional impact, which is not reflected in binary mutation matrices. Fisher’s exact tests were not performed because they would not assess this functional dimension.

**Figure 3:**
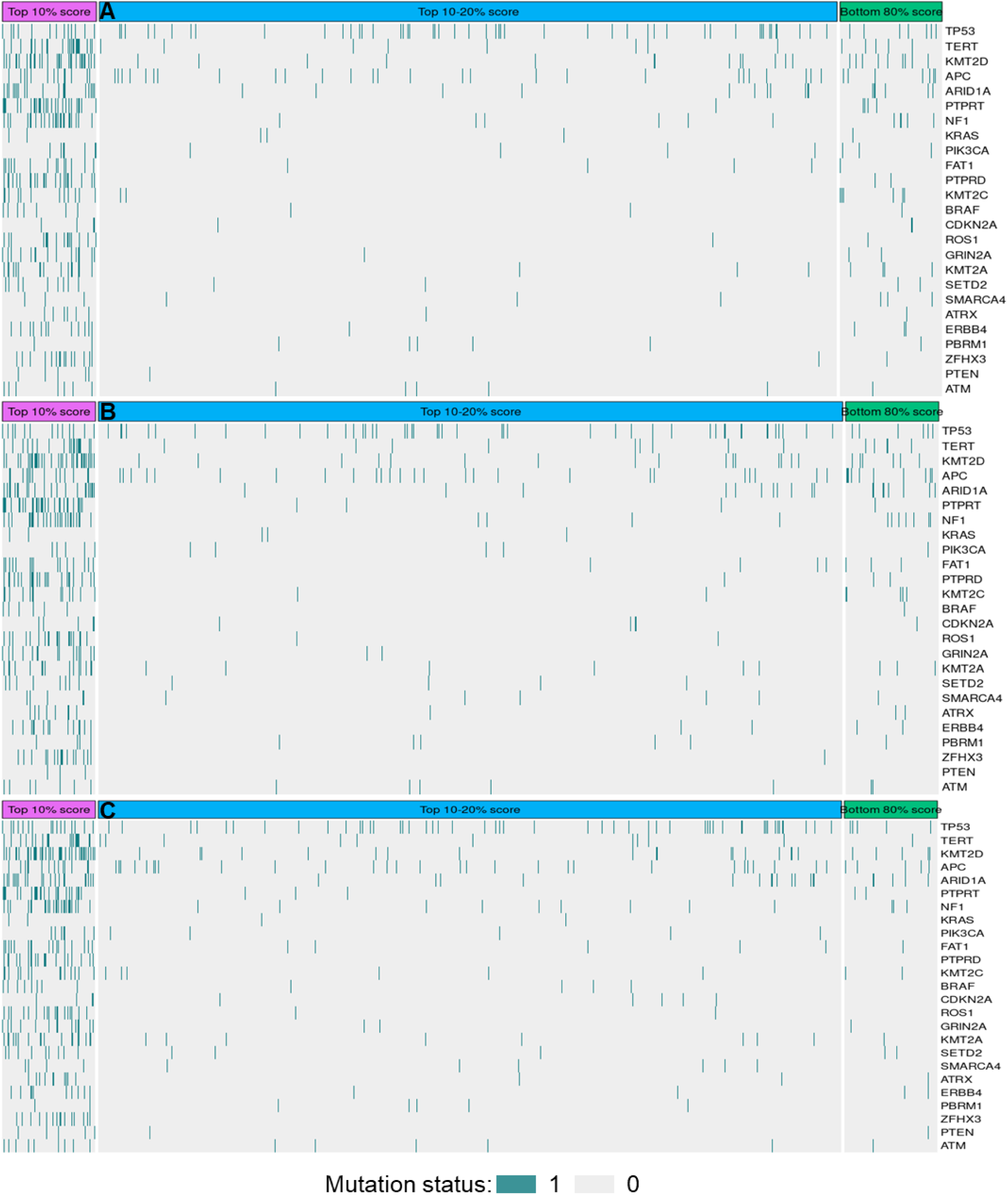
Heatmaps of the Gene Mutation Status provides insights into the distribution of mutational events across different classes of the metrics including (A) Heatmap of gene mutation frequencies stratified by TMB percentile groups (Top 10%, Top 10–20%, Bottom 80%). Higher TMB groups show enriched mutational density across several genes, including *TP53*, *KMT2D*, *TERT*, and *APC*. (B) Heatmap of mutation status grouped by AlphaMissense score classifications. Patterns are similar to TMB but highlight more selective enrichment of functionally significant mutations, with higher Alpha groups exhibiting concentrated alterations in mismatch repair genes and chromatin modifiers. (C) Heatmap stratified by AlphaTMB scores reveals a more focused clustering of biologically relevant mutations, reflecting tumors with both high burden and high predicted pathogenicity. AlphaTMB-high samples cluster mutations in canonical drivers and genes implicated in ICI response; heatmaps depict presence/absence, not functional weight.

**Figure 4.**
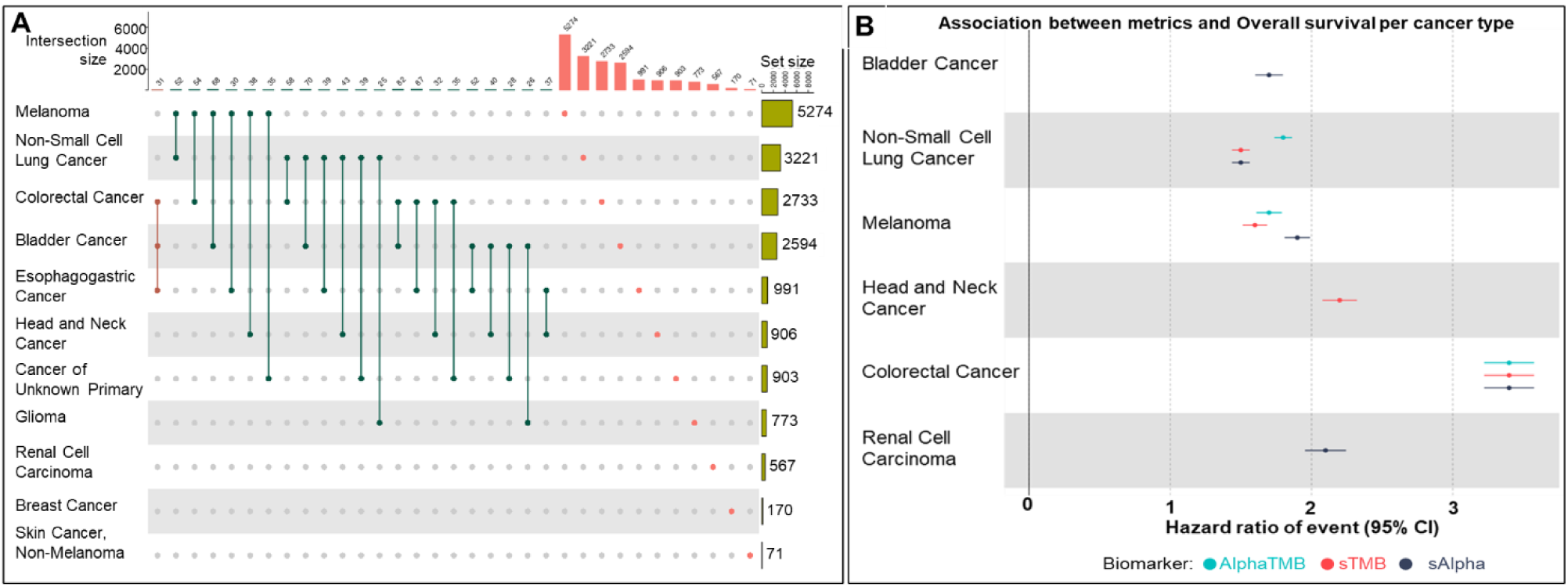
Mutation Co-Occurrence and Biomarker Associations with Survival Across Cancer Types. (A) UpSet plot displaying co-occurring somatic mutations across multiple cancer types in the cohort. Frequent overlaps are observed between melanoma, bladder, and colorectal cancers, suggesting shared oncogenic pathways and immunogenic potential in AlphaMissense-high or TMB-high tumors. (B) Forest plot summarizing hazard ratios for overall survival associated with TMB, AlphaMissense, and AlphaTMB scores across different cancer types. AlphaTMB (cyan) shows comparable or superior prognostic value relative to TMB (red) and Alpha (dark blue) in several tumor types, including melanoma, NSCLC, bladder cancer, and renal cell carcinoma. These findings support the clinical relevance of pathogenicity-weighted mutation burden in stratifying immunotherapy outcomes across diverse malignancies.

### Survival Impact of TMB, AlphaMissense, and AlphaTMB Stratification Using Global and Cancer-Specific Cutoffs

To assess the prognostic utility of TMB, AlphaMissense, and AlphaTMB, we performed multivariable Cox proportional hazards regression analyses using both global and cancer-specific percentile stratifications. As shown in Table 2, which applies pan-cancer thresholds, patients in the Bottom 80% TMB group had a significantly higher risk of death compared to those in the Top 10% group (HR = 2.71, 95% CI: 1.97–3.71, *p* < 0.001). Similar trends were observed for the Alpha metric (HR = 2.62, 95% CI: 1.92–3.59, *p* < 0.001) and AlphaTMB (HR = 2.51, 95% CI: 1.83–3.43, *p* < 0.001), confirming that lower scores across all three metrics are associated with poorer survival. Table 3 presents the same analysis using cancer-type–specific percentiles to account for inter-tumor heterogeneity in mutation burden. While hazard ratios remained statistically significant, effect sizes were slightly attenuated across metrics. This adjustment reduced variability and improved model robustness for tumor-specific comparisons. Importantly, AlphaTMB maintained strong prognostic discrimination, particularly between the Top 10% and Bottom 80% groups, supporting its consistency across different classification strategies. Together, these findings demonstrate that AlphaMissense-informed metrics, especially AlphaTMB, offer prognostic value comparable to or better than traditional TMB, and that their predictive performance is robust across global and tumor-specific stratifications.

**Table 2:** Cox Proportional Hazards Model Using Global Percentile Cutoffs for TMB, Alpha, and AlphaTMB.

| Characteristic | TMB |  |  | Alpha |  |  | AlphaTMB |  |  |
| --- | --- | --- | --- | --- | --- | --- | --- | --- | --- |
|  | HR<sup>1</sup> | 95% CI<sup>1</sup> | p-value | HR<sup>1</sup> | 95% CI<sup>1</sup> | p-value | HR<sup>1</sup> | 95% CI<sup>1</sup> | p-value |
| class_sTMB |  |  |  |  |  |  |  |  |  |
| Top 10% score | — | — |  |  |  |  |  |  |  |
| Bottom 80% score | 2.71 | 1.97, 3.71 | <0.001 |  |  |  |  |  |  |
| Top 10-20% score | 1.51 | 1.02, 2.25 | 0.039 |  |  |  |  |  |  |
| class_sAlpha |  |  |  |  |  |  |  |  |  |
| Top 10% score |  |  |  | — | — |  |  |  |  |
| Bottom 80% score |  |  |  | 2.62 | 1.92, 3.59 | <0.001 |  |  |  |
| Top 10-20% score |  |  |  | 1.54 | 1.03, 2.29 | 0.036 |  |  |  |
| class_AlphaTMB |  |  |  |  |  |  |  |  |  |
| Top 10% score |  |  |  |  |  |  | — | — |  |
| Bottom 80% score |  |  |  |  |  |  | 2.51 | 1.83, 3.43 | <0.001 |
| Top 10-20% score |  |  |  |  |  |  | 2.11 | 1.45, 3.08 | <0.001 |
| <sup>1</sup> HR = Hazard Ratio, CI = Confidence Interval |  |  |  |  |  |  |  |  |  |

Hazard ratios (HRs), confidence intervals (CIs), and p-values for survival analysis across global classification thresholds (Top 10%, Top 10–20%, Bottom 80%) using TMB, AlphaMissense (Alpha), and the composite AlphaTMB. The Top 10% group is used as the reference. AlphaTMB shows improved risk stratification compared to TMB alone. Note: sTMB refers to global percentile-based TMB stratification across the entire cohort, while gTMB refers to stratification within specific cancer group.

**Table 3:** Cox Proportional Hazards Model Using Cancer-Type Specific Percentile Cutoffs.

| Characteristic | TMB |  |  | Alpha |  |  | AlphaTMB |  |  |
| --- | --- | --- | --- | --- | --- | --- | --- | --- | --- |
|  | HR<sup>1</sup> | 95% CI<sup>1</sup> | p-value | HR<sup>1</sup> | 95% CI<sup>1</sup> | p-value | HR<sup>1</sup> | 95% CI<sup>1</sup> | p-value |
| class_gTMB |  |  |  |  |  |  |  |  |  |
| Top 10% score | — | — |  |  |  |  |  |  |  |
| Bottom 80% score | 2.23 | 1.65, 3.02 | <0.001 |  |  |  |  |  |  |
| Top 10-20% score | 1.74 | 1.21, 2.51 | 0.003 |  |  |  |  |  |  |
| class_gAlpha |  |  |  |  |  |  |  |  |  |
| Top 10% score |  |  |  | — | — |  |  |  |  |
| Bottom 80% score |  |  |  | 2.31 | 1.70, 3.13 | <0.001 |  |  |  |
| Top 10-20% score |  |  |  | 1.88 | 1.29, 2.74 | <0.001 |  |  |  |
| class_gAlphaTMB |  |  |  |  |  |  |  |  |  |
| Top 10% score |  |  |  |  |  |  | — | — |  |
| Bottom 80% score |  |  |  |  |  |  | 2.22 | 1.64, 3.00 | <0.001 |
| Top 10-20% score |  |  |  |  |  |  | 1.82 | 1.26, 2.63 | 0.002 |
| <sup>1</sup> HR = Hazard Ratio, CI = Confidence Interval |  |  |  |  |  |  |  |  |  |

Survival associations using stratification based on cancer-specific quantiles for TMB, Alpha, and AlphaTMB. Compared to global cutoffs, this approach reduces inter-tumor variability. AlphaTMB maintains superior prognostic performance, particularly in differentiating the Bottom 80% from the Top 10% group. Note: Note: sTMB refers to global percentile-based TMB stratification across the entire cohort, while gTMB refers to stratification within specific cancer group.

### Mutation Co-Occurrence and Biomarker-Specific Survival Associations Across Cancer Types

To further explore the biological underpinnings and cancer-type–specific relevance of the scoring metrics, we analyzed patterns of co-occurring mutations and the survival impact of TMB, AlphaMissense, and AlphaTMB across distinct tumor types. Figure 3A shows an UpSet plot summarizing recurrent mutation overlaps across the most represented cancers in the cohort. Melanoma and non-small cell lung cancer (NSCLC) exhibited the highest total mutation counts, with shared alterations in genes such as *TP53*, *TERT*, *KMT2D*, and *ARID1A*. These overlaps were particularly prominent in tumors previously classified as high scoring by AlphaMissense or AlphaTMB. In contrast, cancers such as glioma and breast cancer displayed more tumor-specific mutation profiles with fewer shared events. Importantly, tumors harboring co-mutations in mismatch repair genes or chromatin regulators frequently appeared in the top AlphaMissense and AlphaTMB tiers, reinforcing the link between genomic instability and predicted immunotherapy responsiveness.

Figure 3B illustrates the association of each biomarker TMB, AlphaMissense, and AlphaTMB with overall survival across individual cancer types. While all three metrics showed significant associations with survival in multiple cancers, AlphaTMB consistently demonstrated equal or superior prognostic power. For example, AlphaTMB showed comparable prognostic value and aided reclassification in borderline strata including TMB in renal cell carcinoma, bladder cancer, and colorectal cancer. In contrast, TMB showed slightly better performance in head and neck cancer. These results underscore the cancer-type–specific utility of AlphaMissense-informed metrics and suggest that AlphaTMB may offer broader predictive generalizability across tumor types than mutation burden alone. While AlphaTMB showed comparable or directionally favorable performance in certain cancer types, these findings should be interpreted cautiously and validated in larger, tumor-specific cohorts.

## Discussion

This study investigates the prognostic utility of AlphaTMB, a composite biomarker that combines Tumor Mutational Burden (TMB) with AlphaMissense pathogenicity scores, in predicting immunotherapy outcomes across a diverse cohort of cancer patients. By refining mutation burden with information on functional impact, AlphaTMB addresses the limitations of existing biomarkers and provides a biologically enriched predictor of immune checkpoint inhibitor (ICI) response. Pathogenic missense mutations can generate higher quality neoantigens and are often clonal, increasing immune visibility. AlphaTMB integrates mutation quantity and predicted functional impact, enriching tumors with both high burden and immunogenic potential. Pathogenic or driver mutations may better predict response to immune checkpoint inhibitors (ICIs) than total tumor mutational burden (TMB) because they generate high-quality neoantigens and influence tumor–immune interactions. Missense mutations often create novel peptides presented on MHC molecules, which are not subject to central tolerance and can elicit strong T-cell responses [25]. These neoantigens are typically clonal and maintained across tumor cells, enhancing immune visibility [26]. In contrast, passenger mutations often yield subclonal or poorly expressed antigens with limited immunogenicity. Additionally, driver mutations can shape the tumor microenvironment (TME): for example, STK11 and KEAP1 loss-of-function mutations are linked to immune exclusion and ICI resistance, whereas TP53 alterations may promote immune infiltration [27, 28]. These mechanisms support the rationale for integrating mutation burden with predicted functional impact, as implemented in AlphaTMB [29, 30].

While this study did not directly analyze TME features, prior work links mutational burden to T-cell infiltration, checkpoint expression, and HLA diversity [29, 30]. Future studies integrating AlphaTMB with immune profiling and HLA typing could clarify these relationships.

The integration of tumor mutation burden (TMB) into immunotherapy decision-making has marked a significant advance in precision oncology. However, its limitations, particularly its inability to distinguish between biologically meaningful and neutral mutations, have led to inconsistent clinical outcomes and questioned its utility as a standalone biomarker [31–34]. In this study, we addressed these challenges by incorporating a pathogenicity-aware scoring model, AlphaMissense, into TMB calculations to develop AlphaTMB, a composite biomarker that reflects both the quantity and the predicted functional relevance of somatic mutations. Our findings demonstrate that AlphaTMB improves upon traditional TMB-based stratification in predicting survival outcomes among patients treated with immune checkpoint inhibitors (ICIs). Consistent with prior studies that validated TMB as a predictor of ICI responsiveness [35–37], our results reaffirm that higher mutation load correlates with improved survival. However, by integrating AlphaMissense scores, we were able to refine this association and reveal additional layers of prognostic value. Our data collectively provide foundational insights into the relationships between tumor mutational burden (TMB), AlphaMissense scores, and the newly proposed composite metric AlphaTMB, with particular attention to their prognostic relevance in immunotherapy-treated patients. AlphaTMB is not proposed as a definitive benchmark but as an exploratory composite metric that integrates mutation quantity (TMB) with predicted functional impact (AlphaMissense). This design reflects the hypothesis that tumors with both high mutation burden and high pathogenicity scores may exhibit greater immunogenicity. While AlphaTMB demonstrated prognostic discrimination comparable to or slightly improved over TMB in this cohort, its clinical utility remains investigational and requires validation in independent datasets and prospective studies before adoption as a standard biomarker.

The efficacy of ICIs varies widely, prompting a search for robust biomarkers to predict which patients will benefit. Tumor mutational burden (TMB), defined as the number of somatic mutations per megabase, emerged as a key genomic biomarker because a higher mutation load increases the chances of neo-antigen generation and T-cell recognition [38]. Retrospective studies across multiple tumor types have shown that patients with TMB-high tumors respond better to ICIs [39, 40]. An exploratory analysis of the KEYNOTE-158 trial contributed to the U.S. FDA’s approval of pembrolizumab for solid tumors with TMB ≥10 mutations/Mb [41], establishing TMB-high as a tissue-agnostic biomarker for patient selection. However, this broad approval belies important nuances. Only ∼29% of TMB-high patients in that study achieved objective responses, underscoring that high TMB is not synonymous with benefit for all. Subsequent analyses and reviews have raised concerns that a universal TMB cutoff is overly simplistic optimal thresholds likely vary by tumor histology, and TMB may be a surrogate for more direct immunogenic features [42]. For example, microsatellite instability (MSI), resulting from mismatch-repair deficiency, invariably leads to high TMB and is a well-established biomarker of response to ICIs [43]. Similarly, tumors harboring **POLE** mutations exhibit ultrahigh TMB and strong ICI responsiveness due to a hypermutator phenotype, as demonstrated in prior genomic studies [44]. The evolving landscape of immuno-oncology biomarkers suggests that while TMB captures an important dimension of tumor “foreignness,” it often needs refinement or complementation by other factors for precision oncology. In this context, our findings explore AlphaMissense and AlphaTMB as refined biomarker approaches to improve upon TMB alone.

The clinical and demographic characteristics of the 1,529-patient cohort showed that non-small cell lung cancer (NSCLC) and melanoma were the most prevalent cancers, reflecting current clinical patterns where ICIs have demonstrated robust efficacy [45, 46]. The cohort showed a male predominance (63%) and was largely composed of patients aged 50 to 70 years, aligning with global cancer incidence trends and eligibility criteria for immunotherapy [47]. The balanced distribution of survival status (49% deceased vs. 51% living) establishes a well-powered dataset for evaluating genomic predictors of survival and response to treatment.

We then explores the foundational relationships between TMB, AlphaMissense, and patient survival (Figure 1). Figure 1A shows that approximately 69.4% of nonsynonymous variants were annotated with AlphaMissense scores, affirming the broad applicability of this deep learning-based tool in real-world genomic datasets. The strong positive correlation between TMB and AlphaMissense scores (Spearman ρ = 0.866, p < 0.001, Figure 1B) highlights the expected collinearity between the two measures, as tumors with higher mutation loads also tend to harbor more missense variants. This high correlation is expected, given that the included AlphaMissense variants are a subset of total mutations tumors with more mutations generally also have more high-impact missense variants. However, this correlation also obscures the underlying biological diversity between tumors with similar TMB but differing mutation quality. Biologically, AlphaMissense could add nuance by emphasizing the quality of mutations over sheer quantity. For two tumors with equal TMB, the one with a higher AlphaMissense Score harbors a greater proportion of deleterious mutations, which might translate to a higher burden of truly immunogenic neoantigens or more disrupted tumor suppressors. These subtleties are not apparent from TMB alone. In other words, AlphaMissense may act as a lens that reveals differences in the nature of the mutational landscape distinguishing tumors dominated by potentially immunogenic, structural variations from those with mostly synonymous changes. While the strong correlation means AlphaMissense will rarely contradict TMB on a broad scale, our analysis suggests it can refine patient stratification (as seen in the survival outcomes). This implies that AlphaMissense adds biological signals by filtering out mutations less likely to contribute to anti-tumor immunity, thereby complementing TMB. These results echo the sentiment that TMB is often a surrogate measure and that drilling down to more directly relevant genomic features could improve predictive accuracy. Importantly, the density distribution plot indicated that TMB and AlphaMissense are heavily right-skewed distributions, reflecting that most patients have relatively low mutation loads while a minority have extremely high burdens (Figure 1C and 1D). When stratified by survival status, clear distributional differences emerged. Patients who experienced longer survival tends to have higher TMB and AlphaMissense values, on average, compared to those with poorer survival. The density curves for the surviving group were shifted toward the right and showed a longer tail of ultra-high values. In contrast, non-survivors were enriched among patients with low mutation burdens, and their distribution tapered off quickly with far fewer high-TMB outliers. These patterns highlight the prognostic implications of mutation burden: higher mutational load is associated with improved patient outcomes under ICI therapy, as supported by multiple large-scale analyses [24, 48, 49] 2019). This pattern supports the hypothesis that not only the quantity, but the functional quality of mutations, specifically their capacity to produce immunogenic neoantigens plays a key role in therapeutic success [31].

We further introduced AlphaTMB as a hybrid biomarker that integrates both TMB and AlphaMissense information, intending to refine patient classification beyond what either metric provides alone. (Figure 2). In practice, AlphaTMB reassigns patients by considering how many mutations their tumor has and how many of those mutations are of high functional significance. This composite approach improved classification accuracy in our analyses effectively enhancing the identification of true high-risk or low-risk patients. For instance, some patients who were borderline TMB-high by count were found to have relatively low AlphaMissense (i.e. their mutations were mostly benign); AlphaTMB downgraded such cases, which correlated with poorer outcomes than the raw TMB-high group. Conversely, a few patients with intermediate TMB but disproportionately high AlphaMissense Score were upgraded by AlphaTMB, identifying cases of modest mutation burden where an unusually large fraction of mutations were deleterious (and who indeed had outcomes more like the TMB-high group). By reassigning these discrepant cases, AlphaTMB achieved a cleaner separation of patients in survival analyses on the global scale, suggesting it is a more precise pan cancer classifier of immunotherapy benefit. This outcome resonates with the broader notion in precision oncology that composite biomarkers outperform single metrics [50]. Others have advocated combining TMB with additional variables such as neoantigen quality, immune gene expression, or HLA genotype to improve predictive power [21, 51, 52]. AlphaTMB exemplifies this strategy by fusing mutation quantity and quality. Clinically, such a hybrid metric could be valuable. It might be used to refine eligibility criteria or risk stratification: for example, two patients above a TMB cutoff might be differentiated by AlphaTMB, flagging one as a likely responder and the other as questionable. This reinforces the added discriminatory power of AlphaTMB in resolving cases where TMB alone is ambiguous, a challenge well-documented in the literature [53, 54].

The gene mutation heatmaps (Figure 3) provide insight into the genomic landscape of high-scoring tumors. We observed that certain mutations clustered within the high TMB and AlphaMissense groups, pointing to specific biological subsets enriched for neo-antigen load. Notably, DNA mismatch repair (MMR) gene mutations (such as *KMT2D, TERT, PTPRT,* or *ARID1A*) were concentrated in the tumors with top-tier TMB/AlphaMissense scores. This is expected, as loss-of-function in MMR genes causes microsatellite instability and hypermutation; tumors with these alterations accrue hundreds of mutations and have demonstrated high response rates to PD-1 blockade. Their presence in the high-score cluster validates that our metrics capture these hypermutated, immunotherapy-sensitive cases. Their clustering in the heatmap further supports AlphaMissense/AlphaTMB as capturing “neoantigen-rich” genotypes. We also noted that common driver mutations like TP53 and KRAS were prevalent across many high-scoring tumors. While TP53 mutations are ubiquitous in many cancers regardless of TMB, their co-occurrence with high AlphaMissense indicates tumors with both driver events and a high load of other deleterious mutations. In some cases, specific co-mutation patterns emerged: for example, in lung cancers with high scores, *KRAS* mutations often co-occurred with others and have been linked to an inflammatory tumor microenvironment conducive to ICI response. These heatmap observations imply that high AlphaMissense or AlphaTMB groups are not random collections of mutations, they are enriched for genomic alterations (MMR deficiency, POLE mutation, certain oncogenic drivers) that biologically underpin a high neoantigen load and immunoresponsive phenotype. This concordance between specific mutation clusters and our scoring groups lends mechanistic credibility to AlphaMissense: it picks known genomic hallmarks of immunotherapy-responsive tumors rather than just arbitrary mutation counts.

The UpSet plot further corroborates the observed pattern from the heatmap. The intersection of mutation sets across different cancer types showed that the AlphaMissense captures biologically meaningful co-mutation patterns. Indeed, the analysis revealed that certain combinations of mutations occur across multiple tumor types, particularly among high AlphaMissense tumors. For example, a set of mismatches repair gene mutations (such as *KMT2D* and *TERT* together) was found in AlphaMissense-high cases of colorectal and endometrial cancers, suggesting the cross-cancer occurrence of MMR gene mutations. This co-occurrence is biologically meaningful: these cancers, despite different tissue origin, share the MMR-deficient, hypermutated condition that confers sensitivity to PD-1 inhibitors. AlphaMissense successfully flagged these cases by their high load of damaging mutations. The biomarker thus captures the commonality that these tumors all have extreme mutation loads and likely high neoantigen counts, even though they arise in different organs. We also observed co-occurring driver mutation patterns in AlphaMissense-high groups across cancers: for instance, the combination of *TP53* and *KRAS* mutations appears in both lung adenocarcinoma and pancreatic cancer subsets. Tumors harboring both TP53 and KRAS mutations have aggressive biology but also tend to exhibit genomic instability; in NSCLC, KRAS-mutant tumors often have inflamed microenvironments that could favor ICI response. Such overlaps in the UpSet plot indicate that AlphaMissense is sensitive to mutation patterns that transcend tumor type, many known to modulate immunotherapy outcomes.

The Cox proportional hazards modeling of overall survival further highlights the prognostic relevance of TMB, AlphaMissense, and AlphaTMB. In our analysis, all three biomarkers showed significant associations with patient survival on immunotherapy, but with varying degrees of effect.

To assess model stability and generalizability, we performed bootstrap validation with 100 iterations. In each iteration, training data were generated by sampling with replacement from the original dataset, while out-of-bag samples served as the independent test set Supplementary table1-3.High TMB was associated with improved survival (reduced hazard of death) relative to low TMB, which aligns with prior multivariate studies showing that TMB-high patients often have longer median survival under ICIs. AlphaMissense yielded a similar hazard trend, confirming that a high load of deleterious missense mutations correlates with better survival. Importantly, the AlphaTMB composite had the strongest hazard ratio effect among the three. Patients classified as high by AlphaTMB had the most pronounced survival advantage. In contrast, those flagged as low by AlphaTMB fared the worst, with the model indicating a wider separation between risk groups than TMB or AlphaMissense alone. This suggests that AlphaTMB discriminates more in prognostication. One interpretation is that by integrating two layers of information, AlphaTMB identifies a subset of patients who are *truly* likely to benefit (hence pulling their survival curve up) and filters out some who would not (pushing the non-responder curve down). In practical terms, the composite biomarker captured the survival signal of TMB. Still, it sharpened it by removing cases where TMB was high in quantity but low in quality, and vice versa. These findings resonate with the emerging consensus that multi-parametric biomarkers provide more robust prognostic information in immunotherapy. It is also noteworthy that TMB and AlphaMissense are not entirely independent. Although TMB and AlphaMissense are strongly correlated, they capture different aspects of tumor biology. TMB measures mutation quantity, while AlphaMissense estimates the functional impact of missense variants. Tumors with similar TMB can differ in the pathogenicity of their mutations, which may influence immunogenicity. AlphaTMB combines these dimensions by weighting mutation burden with predicted deleteriousness, thereby prioritizing tumors enriched for functionally significant alterations. This approach may improve patient stratification, particularly in borderline TMB cases, and aligns with the principle that both mutation load and biological relevance shape immune checkpoint inhibitor response.

Our analysis compares three related biomarkers TMB, AlphaMissense, and the hybrid AlphaTMB and demonstrates their relative strengths in prognosticating immunotherapy outcomes. TMB is an established metric with proven but imperfect predictive value. It provides a broad measure of tumor mutation quantity and has been validated in retrospective studies and clinical trials as an indicator of response, leading to its clinical adoption. However, TMB alone can be noisy; tumor types differ in what constitutes “high” TMB, and many TMB-high patients do not benefit from ICIs while some TMB-low patients do. AlphaMissense represents a refinement by focusing on the qualitative subset of mutations most likely to be biologically impactful, it hones the TMB signal. We found that AlphaMissense strongly correlates with TMB (indicating it doesn’t discard the broad signal). Yet, it improved risk stratification and aligned with known immunogenic patterns more closely than TMB. This suggests that AlphaMissense could serve as a more specific surrogate for neo-antigen load, filtering out passengers and emphasizing biologically meaningful mutations. AlphaTMB, in turn, combines the best of both: it uses the comprehensive scope of TMB but tempers it with the specificity of AlphaMissense. Our results show that AlphaTMB had the highest prognostic accuracy, indicating that a composite approach can add value beyond either metric alone which is a finding that accentuates the for integrated biomarker models. Technically, all three metrics are interrelated (as evidenced by high TMB–AlphaMissense correlation), yet those subtle differences that AlphaMissense captures make a tangible difference when formalized into AlphaTMB. The development of robust biomarkers for immunotherapy requires integrating genomic predictors with tumor microenvironment (TME) features and functional immune readouts. While AlphaTMB focuses on pathogenicity-weighted mutation burden, its clinical utility will be enhanced by combining it with transcriptomic, proteomic, and immune profiling data. Advances in multi-omics technologies and computational drug discovery frameworks provide methodological pathways for translating AlphaTMB into clinical decision-making [55, 56]. Additionally, in-depth research on immunotherapy mechanisms and molecular pharmacology offers theoretical support for understanding the biological basis of AlphaTMB’s predictive performance [57–59]. Future studies should explore these integrative approaches to refine patient selection and optimize therapeutic outcomes.

### Limitations

This study has several limitations. First, AlphaMissense currently predicts the pathogenicity of missense variants only, excluding other immunogenically relevant alterations such as frameshift insertions/deletions, nonsense mutations, and non-coding variants that can also generate neoantigens. Prediction accuracy for rare variants and applicability across diverse populations require further evaluation to minimize bias. Second, AlphaTMB thresholds may require tumor-type–specific calibration for optimal discrimination. Although AlphaMissense scores were precomputed for approximately 71 million variants—making AlphaTMB calculation computationally trivial—clinical implementation will necessitate harmonization with existing biomarker reporting standards and validation in prospective, multi-platform cohorts. Third, this analysis is based on a single retrospective cohort (MSK-IMPACT), which limits external generalizability. Future validation should leverage independent ICI-treated cohorts, such as prospective clinical trials or real-world registries, with harmonized sequencing and treatment annotation. Technical harmonization should include panel normalization, variant-calling consistency, and alignment with ongoing TMB harmonization efforts. While AlphaTMB is platform-agnostic because it relies on precomputed AlphaMissense scores and normalized mutation counts, cross-platform benchmarking and tumor-specific thresholds remain essential for clinical translation. Finally, clinical covariates such as ECOG performance status, PD-L1 expression, MSI/dMMR status, HLA genotype, and tumor microenvironment features were unavailable in this dataset. Future work should validate AlphaTMB in independent cohorts, benchmark it against established biomarkers, and integrate it with transcriptomic and immune profiling approaches (e.g., IOBR, GseaVis) to contextualize its predictive value within the tumor microenvironment.

## CONCLUSION

This study presents AlphaTMB as a novel and clinically relevant biomarker that integrates tumor mutational burden (TMB) with AlphaMissense pathogenicity scores to enhance prediction of response to immune checkpoint inhibitors. By combining the quantity of mutations with their predicted functional impact, AlphaTMB offers a more refined and biologically meaningful measure of a tumor’s immunogenic potential compared to TMB alone. The findings demonstrate that AlphaTMB consistently outperforms both TMB and AlphaMissense scores individually in stratifying patients by survival outcomes, with comparable prognostic value and improved reclassification of borderline risk groups. A key takeaway from this research is that pathogenicity-informed metrics can provide a higher-resolution view of tumor biology and are better aligned with the mechanisms underlying immunotherapy response. The enrichment of mismatch repair and chromatin remodeling gene mutations in AlphaTMB-high tumors supports its ability to capture biologically relevant, neoantigen-rich phenotypes associated with favorable immune engagement. However, the study also acknowledges several limitations. First, although AlphaMissense scores provide significant insight into missense variants, the model does not yet account for other immunogenically important mutation types such as frameshift insertions/deletions, nonsense mutations, or non-coding variants that can also generate neoantigens. Second, this work is based on retrospective data from a single cohort (MSK-IMPACT), and while the results are promising, prospective validation across independent datasets and cancer types is essential to confirm generalizability. Third, while AlphaTMB showed pan-cancer applicability, tumor-type–specific thresholds may further enhance its predictive precision and should be explored in future analyses. Looking ahead, future research should focus on expanding the scope of AlphaMissense to incorporate additional variant classes and integrating AlphaTMB into multimodal biomarker frameworks alongside transcriptomic, epigenetic, or immunological data. Additionally, clinical studies are needed to assess the utility of AlphaTMB in real-time decision-making, particularly in guiding therapy for patients with intermediate or ambiguous TMB profiles. Ultimately, the adoption of AlphaTMB could mark a meaningful advance in the personalization of immunotherapy, offering more accurate patient selection and potentially improving treatment outcomes in diverse cancer settings.

**Future studies should integrate AlphaTMB with other biomarkers such as PD-L1 expression, MSI/dMMR status, and treatment line to comprehensively assess clinical utility. Additionally, incorporating transcriptomic and immune profiling using tools such as IOBR and GseaVis could contextualize AlphaTMB within the tumor microenvironment and identify mechanistic pathways underlying immunotherapy response.**

## Data Availability

All clinical and mutational data used in this study are publicly available from the MSK-IMPACT cohort via cBioPortal (https://www.cbioportal.org). AlphaMissense variant scores were obtained from the publicly released DeepMind AlphaMissense dataset.

## Ethics statement

This study utilized publicly available data and software-based analysis, which does not involve any kind of collection and distribution of human subject data.

## Appendix

**Table1:** Bootstrap summary.

| Total Iteration | Successful iterations | Success rate | Failed iterations |
| --- | --- | --- | --- |
| 100 | 100 | 1 | 0 |

**Table 2:** Test size summary.

| Mean test size | sd test size | Min test size | Max test size | Median test size |
| --- | --- | --- | --- | --- |
| 564.67 | 11.66 | 519 | 596 | 565.5 |

**Table 3:**
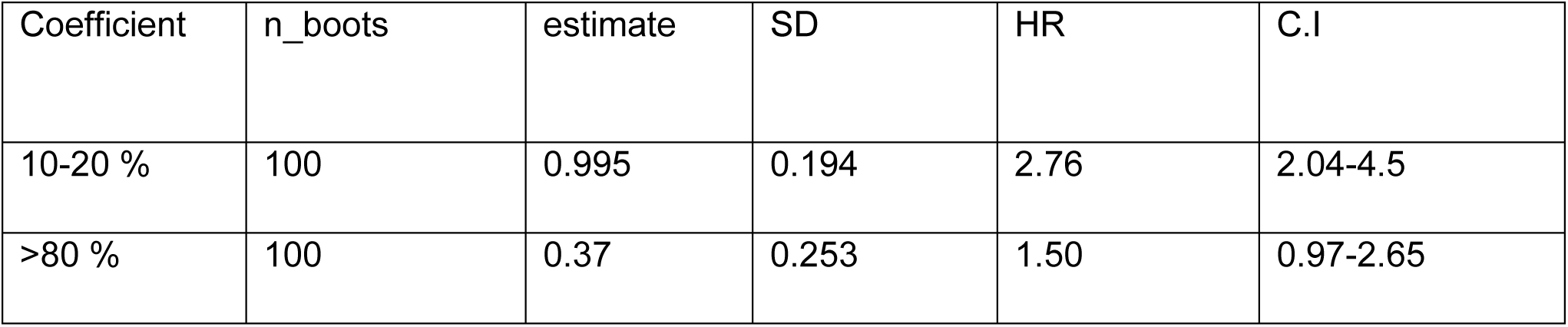
Coefficient stability.

| Coefficient | n_boots | estimate | SD | HR | C.I |
| --- | --- | --- | --- | --- | --- |
| 10-20 % | 100 | 0.995 | 0.194 | 2.76 | 2.04-4.5 |
| >80 % | 100 | 0.37 | 0.253 | 1.50 | 0.97-2.65 |

## References

1. Sung, H., et al., Global cancer burden in 2022: Incidence, mortality and prevalence. CA: A Cancer Journal for Clinicians, 2024.

2. Bray, F., et al., The ever-increasing importance of cancer as a leading cause of premature death worldwide. Cancer, 2021. 127(16): p. 3029–3030.

3. Dare, A.J., et al., High-burden cancers in Middle-income countries: a review of Prevention and early detection strategies targeting At-risk populations. Cancer Prevention Research, 2021. 14(12): p. 1061–1074.

4. De Souza, J.A., et al., Global health equity: cancer care outcome disparities in high-, middle-, and low-income countries. Journal of Clinical Oncology, 2016. 34(1): p. 6–13.

5. Shiravand, Y., et al., Immune checkpoint inhibitors in cancer therapy. Current Oncology, 2022. 29(5): p. 3044–3060.

6. Sharma, P., et al., Immune checkpoint therapy—current perspectives and future directions. Cell, 2023. 186(8): p. 1652–1669.

7. Wei, J., et al., Current trends in sensitizing immune checkpoint inhibitors for cancer treatment. Molecular Cancer, 2024. 23(1): p. 279.

8. Zhang, Y. and Z. Zhang, The history and advances in cancer immunotherapy: understanding the characteristics of tumor-infiltrating immune cells and their therapeutic implications. Cellular & molecular immunology, 2020. 17(8): p. 807–821.

9. Ribas, A. and J.D. Wolchok, Cancer immunotherapy using checkpoint blockade. Science, 2018. 359(6382): p. 1350–1355.

10. Yarchoan, M., et al., Targeting neoantigens to augment antitumour immunity. Nature Reviews Cancer, 2017. 17(4): p. 209–222.

11. Goloudina, A., et al., Shared neoantigens for cancer immunotherapy. Molecular Therapy Oncology, 2025. 33(2).

12. Huang, T., et al., Prognostic role of tumor mutational burden in cancer patients treated with immune checkpoint inhibitors: a systematic review and meta-analysis. Frontiers in oncology, 2021. 11: p. 706652.

13. Fumet, J.-D., et al., Tumour mutational burden as a biomarker for immunotherapy: current data and emerging concepts. European Journal of Cancer, 2020. 131: p. 40–50.

14. Marabelle, A., et al., Association of tumour mutational burden with outcomes in patients with advanced solid tumours treated with pembrolizumab: Prospective biomarker analysis of the multicohort, open-label, phase 2 KEYNOTE-158 study. The Lancet Oncology, 2020. 21(10): p. 1353–1365.

15. Zhang, S., et al., Tumor initiation and early tumorigenesis: molecular mechanisms and interventional targets. Signal transduction and targeted therapy, 2024. 9(1): p. 149.

16. Ragone, C., et al., Lack of shared neoantigens in prevalent mutations in cancer. Journal of Translational Medicine, 2024. 22(1): p. 344.

17. Lakatos, E., et al., Evolutionary dynamics of neoantigens in growing tumors. Nature genetics, 2020. 52(10): p. 1057–1066.

18. Vitale, I., et al., Mutational and antigenic landscape in tumor progression and cancer immunotherapy. Trends in Cell Biology, 2019. 29(5): p. 396–416.

19. Sha, D., et al., Tumor Mutational Burden (TMB) as a Predictiv-e biomarker in solid tumors. Cancer Discov 10: 1808–1825. 2020.

20. Vega, D., et al., Aligning tumor mutational burden (TMB) quantification across diagnostic platforms: phase II of the Friends of Cancer Research TMB Harmonization Project. Annals of Oncology, 2021. 32(12): p. 1626–1636.

21. Ahmed, J., et al., Challenges and Future Directions in the Management of Tumor Mutational Burden-High (TMB-H) Advanced Solid Malignancies. Cancers, 2023. 15(24): p. 5841.

22. Budczies, J., et al., Tumour mutational burden: clinical utility, challenges and emerging improvements. Nature Reviews Clinical Oncology, 2024. 21(10): p. 725–742.

23. Cheng, J., et al., Accurate proteome-wide missense variant effect prediction with AlphaMissense. Science, 2023. 381(6664): p. eadg7492.

24. Samstein, R.M., et al., Tumor mutational load predicts survival after immunotherapy across multiple cancer types. Nature genetics, 2019. 51(2): p. 202–206.

25. Łuksza, M., et al., A neoantigen fitness model predicts tumour response to checkpoint blockade immunotherapy. Nature, 2017. 551(7681): p. 517–520.

26. McGranahan, N., et al., Clonal neoantigens elicit T cell immunoreactivity and sensitivity to immune checkpoint blockade. Science, 2016. 351(6280): p. 1463–1469.

27. Dong, Z.-Y., et al., Potential biomarker for checkpoint blockade immunotherapy and treatment strategy. Tumor Biology, 2016. 37(4): p. 4251–4261.

28. Skoulidis, F., et al., STK11/LKB1 mutations and PD-1 inhibitor resistance in KRAS-mutant lung adenocarcinoma. Cancer discovery, 2018. 8(7): p. 822–835.

29. Litchfield, K., et al., Meta-analysis of tumor-and T cell-intrinsic mechanisms of sensitization to checkpoint inhibition. Cell, 2021. 184(3): p. 596–614. e14.

30. Rosenthal, R., et al., Neoantigen-directed immune escape in lung cancer evolution. Nature, 2019. 567(7749): p. 479-485.

31. Jardim, D.L., et al., The challenges of tumor mutational burden as an immunotherapy biomarker. Cancer cell, 2021. 39(2): p. 154–173.

32. Chan, T.A., et al., Development of tumor mutation burden as an immunotherapy biomarker: utility for the oncology clinic. Annals of Oncology, 2019. 30(1): p. 44–56.

33. Zgura, A., et al., Evaluating Tumour Mutational Burden as a Key Biomarker in Personalized Cancer Immunotherapy: A Pan-Cancer Systematic Review. Cancers, 2025. 17(3): p. 480.

34. Galuppini, F., et al., Tumor mutation burden: from comprehensive mutational screening to the clinic. Cancer cell international, 2019. 19: p. 1–10.

35. Braganca Xavier, C., et al., Identifying predictors of overall survival among patients with TMB-low metastatic cancer treated with immune checkpoint inhibitors. Oncologist, 2025. 30(4).

36. Wang, X., et al., Tumor mutational burden for the prediction of PD-(L) 1 blockade efficacy in cancer: challenges and opportunities. Annals of Oncology, 2024. 35(6): p. 508–522.

37. Strickler, J.H., B.A. Hanks, and M. Khasraw, Tumor mutational burden as a predictor of immunotherapy response: is more always better? Clinical Cancer Research, 2021. 27(5): p. 1236–1241.

38. Wang, P., Y. Chen, and C. Wang, Beyond tumor mutation burden: tumor neoantigen burden as a biomarker for immunotherapy and other types of therapy. Frontiers in oncology, 2021. 11: p. 672677.

39. Killock, D., TMB—a histology-agnostic predictor of the efficacy of ICIs? Nature reviews Clinical oncology, 2020. 17(12): p. 718–718.

40. Palmeri, M., et al., Real-world application of tumor mutational burden-high (TMB-high) and microsatellite instability (MSI) confirms their utility as immunotherapy biomarkers. ESMO open, 2022. 7(1): p. 100336.

41. Marabelle, A., et al., Association of tumour mutational burden with outcomes in patients with advanced solid tumours treated with pembrolizumab: prospective biomarker analysis of the multicohort, open-label, phase 2 KEYNOTE-158 study. The Lancet Oncology, 2020. 21(10): p. 1353–1365.

42. Mo, S.-F., et al., Universal cutoff for tumor mutational burden in predicting the efficacy of anti-PD-(L) 1 therapy for advanced cancers. Frontiers in Cell and Developmental Biology, 2023. 11: p. 1209243.

43. Yang, S.-R., et al., Microsatellite instability and mismatch repair deficiency define a distinct subset of lung cancers characterized by smoking exposure, high tumor mutational burden, and recurrent somatic MLH1 inactivation. Journal of Thoracic Oncology, 2024. 19(3): p. 409–424.

44. Kungulovski, G., et al., Tumors with mutations in chromatin regulators are associated with higher mutational burden and improved response to checkpoint immunotherapy. medRxiv, 2024: p. 2024.10. 15.24315153.

45. Schoenfeld, A.J. and M.D. Hellmann, Acquired resistance to immune checkpoint inhibitors. Cancer cell, 2020. 37(4): p. 443–455.

46. Vogel, J., et al., Prospective assessment of demographic characteristics associated with worse health related quality of life measures following definitive chemoradiation in patients with locally advanced non-small cell lung cancer. Translational Lung Cancer Research, 2019. 8(4): p. 332.

47. Siegel, R.L., A.N. Giaquinto, and A. Jemal, Cancer statistics, 2024. CA: a cancer journal for clinicians, 2024. 74(1): p. 12–49.

48. Kim, J.Y., et al., Tumor mutational burden and efficacy of immune checkpoint inhibitors: a systematic review and meta-analysis. Cancers, 2019. 11(11): p. 1798.

49. Sholl, L.M., et al., The promises and challenges of tumor mutation burden as an immunotherapy biomarker: a perspective from the International Association for the Study of Lung Cancer Pathology Committee. Journal of Thoracic Oncology, 2020. 15(9): p. 1409–1424.

50. La Thangue, N.B. and D.J. Kerr, Predictive biomarkers: a paradigm shift towards personalized cancer medicine. Nature reviews Clinical oncology, 2011. 8(10): p. 587–596.

51. Anzar Martínez de Lagrán, I., Optimizing tumor variant detection and HLA typing for neoantigen prediction in cancer immunotherapy. 2024.

52. Tzenios, N., A meta-analysis of cancer immunotherapy: Evaluating efficacy, predictive biomarkers, and therapeutic resistance. 2022: SR21-Institute for Scientific Research.

53. Schnidrig, D., S. Turajlic, and K. Litchfield, Tumour mutational burden: primary versus metastatic tissue creates systematic bias. Immuno-oncology technology, 2019. 4: p. 8–14.

54. McGrail, D., et al., High tumor mutation burden fails to predict immune checkpoint blockade response across all cancer types. Annals of Oncology, 2021. 32(5): p. 661–672.

55. Wang, Z., Y. Zhao, and L. Zhang, Emerging trends and hot topics in the application of multi-omics in drug discovery: a bibliometric and visualized study. Current Pharmaceutical Analysis, 2024.

56. Liu, Y., et al., Advances in drug discovery based on network pharmacology and omics technology. Current Pharmaceutical Analysis, 2024.

57. Li, G., et al., Breaking boundaries: chronic diseases and the frontiers of immune microenvironments. Med Research, 2025.

58. Abdollahi, E., et al., Immunomodulatory Therapeutic Effects of Curcumin on M1/M2 Macrophage Polarization in Inflammatory Diseases. Curr Mol Pharmacol, 2023. 16(1): p. 2–14.

59. Sensi, B., et al., Mechanism, Potential, and Concerns of Immunotherapy for Hepatocellular Carcinoma and Liver Transplantation. Curr Mol Pharmacol, 2024. 17: p. e18761429310703.

